# Micrografting device for testing environmental conditions for grafting and systemic signaling in Arabidopsis

**DOI:** 10.1101/2019.12.20.885525

**Authors:** Hiroki Tsutsui, Naoki Yanagisawa, Yaichi Kawakatsu, Shuka Ikematsu, Yu Sawai, Ryo Tabata, Hideyuki Arata, Tetsuya Higashiyama, Michitaka Notaguchi

## Abstract

**Summary:** Grafting techniques have been applied in studies of systemic, long-distance signaling in several model plants. Seedling grafting in Arabidopsis, known as micrografting, enables investigation of the molecular mechanisms of systemic signaling between shoots and roots. However, conventional micrografting requires a high level of skill, limiting its use. Thus, an easier user-friendly method is needed. Here, we developed a silicone microscaled device, the micrografting chip, to obviate the need for training and to generate less stressed and more uniformly grafted seedlings. The chip has tandemly arrayed units, each of which consists of a seed pocket for seed germination and a micro-path with pairs of pillars for hypocotyl holding. Grafting, including seed germination, micrografting manipulation, and establishment of tissue reunion, is performed on the chip. Using the micrografting chip, we evaluated the effect of temperature and the carbon source on grafting and showed that a temperature of 27°C and a sucrose concentration of 0.5% were optimal. We also used the chip to investigate the mechanism of systemic signaling of iron status using a quadruple nicotianamine synthase (*nas*) mutant. The constitutive iron-deficiency response in the *nas* mutant because of aberrant partitioning was significantly rescued by grafting of wild-type shoots or roots, suggesting that shoot-and root-ward translocation of nicotianamine–iron complexes is essential for iron mobilization. Thus, our micrografting chip will promote studies of long-distance signaling in plants.

**Significance Statement:** A number of micrografting studies on systemic, long-distance signaling have been performed, but the technique is not yet used widely. Here, we developed a silicone-based micrografting chip to improve the ease-of-use, efficiency, and success rate of micrografting, even for untrained users.

## Introduction

Grafting has long been used in agriculture to control fruit-tree cultivation and increase the production and/or promote disease tolerance of vegetables such as *Cucurbitaceae* and *Solanaceae* (Lee and Oda 2003). Grafting is also used in studies of systemic control of plant physiology and its molecular basis. Plants transmit information cues systemically in response to abiotic and biotic signals and use factors such as phytohormones, proteins, and RNA for organ-to-organ communication. Grafting can be used to investigate the transmissibility of physiological signals between a plant and its grafted partner plant over the graft junction and to evaluate molecular transport as the basis of long-distance signaling. Stem grafting, a commonly-used method, involves grafting an upper stem (known as the scion) onto another stem (the stock plant). Another method, hypocotyl grafting of young seedlings, termed micrografting, has been applied in several model plants, such as petunia and Arabidopsis (Tsutsui and Notaguchi, 2017). Micrografting enables investigation of the early developmental stages. Micrografting is useful especially in Arabidopsis because it has a short life cycle. Turnbull *et al.* (2002) established a micrografting method in Arabidopsis, in which genetic resources are abundant; the method has since been widely used in plant science. Important insights have been gained from micrografting experiments, including identification of systemic signals related to flowering (Corbesier *et al.*, 2007, Notaguchi *et al.*, 2008), the nutrient response (Pant *et al.*, 2008, Tabata *et al.*, 2014), the dehydration response (Takahashi *et al.*, 2018), and gene silencing (Molnar *et al.*, 2010) where I-shaped (one-shoot and one-root) grafting or Y-shaped (two-shoot and one-root) grafting were used to investigate shoot-to-root, root-to-shoot, or shoot-to-shoot signaling (Tsutsui and Notaguchi, 2017). Therefore, micrografting is an important technique for studying systemic signaling in plants.

Grafting experiments have been conducted to identify the molecular species transmitted through vascular tissues as long-distance signals. The phytohormone cytokinins have different side chain types and there is preference for transport in xylem and phloem (Sakakibara. 2006). Osugi *et al.* (2017) conducted micrografting experiments using various cytokinin biosynthetic and transport mutants and revealed that root-to-shoot translocation of *trans*-zeatin-riboside and *trans*-zeatin regulates shoot architecture. Like phytohormones, macro-and micronutrients also have each specific form for long-distance transport (Maathuis, 2009, Hänsch and Mendel, 2009). As an instance, iron (Fe) homeostasis is regulated in a systemic manner, where Fe makes conjugates with metal chelators such as citrate (Ci) and nicotianamine (NA) and distributed to entire body via the vascular tissues together with the action of transporters (Maas *et al.*, 1988, Durrett *et al.*, 2007, Gayomba *et al.*, 2015, Kumar *et al.*, 2017). In Arabidopsis harboring a mutation in *Oligopeptide Transporter 3* (*OPT3*), which mediates uptake of Fe into phloem and regulates shoot-to-root Fe redistribution, shoot-to-root Fe transport was disrupted, causing accumulation of Fe in shoots (Zhai *et al.*, 2014). Moreover, the roots of an *opt3* mutant of Arabidopsis exhibited constitutive Fe-deficiency transcriptional responses (Stacey *et al.*, 2008), which were rescued by the grafting of wild-type (WT) shoots (Zhai *et al.*, 2014). Similar to the *opt3* mutant, loss-of-function mutations in nicotianamine synthase (*NAS*) in Arabidopsis and tomato resulted in Fe accumulation in shoots and constitutive Fe-deficiency responses in roots (Scholz *et al.*, 1985, Klatte *et al.*, 2009). NAS is an enzyme that catalyzes the trimerization of *S*-adenosylmethionine to synthesize NA (Ling *et al.*, 1999). NA facilitates the lateral distribution of Fe from xylem to neighboring cells as well as its transport from phloem to sink organs (DiDonato *et al.*, 2004; Schuler *et al.*, 2012). Schuler *et al.* (2012) showed that the chlorotic phenotype of the shoot of an Arabidopsis quadruple mutant in four *NAS* genes, *nas4x-2*, was rescued by NA application directly onto the leaves, providing evidence of NA local activity in Fe homeostasis. However, whether root- or shoot-synthesized NA contributes to Fe homeostasis at the whole-plant level is unclear.

Although the micrografting technique has enabled investigation of the systemic role of genes and of molecular transport, a high level of skill is required. In our experience, the conventional micrografting method requires much practice to achieve a high success rate. Several papers have reported modifications that increase the success rate of micrografting by modifying equipment and procedures (Notaguchi *et al.*, 2009), using an agar-based support (Marsch-Martínez *et al.*, 2013), and using a fine pin to fix plants in position (Huang and Yu, 2015). Nevertheless, micrografting is difficult, particularly for beginners, which hampers its adoption. To improve accuracy and achieve high-throughput analysis, a new concept of engineering techniques to manipulate organisms are required. Micro Electro Mechanical Systems (MEMS), are suitable for microscale biological experiments under controlled conditions. A few case studies have applied these techniques in plants. For example, root growth was accomplished in a microfluidic device designed for cultivation and imaging (Grossmann *et al.*, 2011). Microcages were useful to fix positions and culture plant ovules for long-term observation (Park *et al.*, 2014). Also, tip growth in plant cells, such as pollen tubes, root hairs, and moss protonemata, was tested in specialized devices with narrow gaps (Yanagisawa *et al.*, 2017). Thus, micro-engineering systems enable manipulation of plants. Because a user-friendly method is needed, we applied microengineering techniques to improve micrografting.

Here, we developed a device made of silicone rubber for Arabidopsis micrografting, the micrografting chip, which was fabricated by MEMS technology using poly(dimethylsiloxane) (PDMS). The chip facilitated micrografting and yielded uniformly grafted plants. We demonstrated the utility of the chip by testing the effect of grafting conditions. Finally, the results of grafting experiments using a mutant of *NAS* (involved in long-distance Fe transport) provided insight into systemic Fe homeostasis.

## Results

### The micrografting chip

We developed a micrografting chip (approximately 3.6 × 17 mm) composed of the silicon elastomer PDMS to perform seedling micrografting in Arabidopsis (Figure 1a). One chip contains four in-line units and has a cover (Figure 1b), which is placed on the side of the chip (Figure 1c, d). Each unit consists of a seed pocket for sowing seeds, spaces for root growth below the seed pocket, and a micro-path with pairs of micro-pillars for setting a hypocotyl above the seed pocket. A horizontal slot across the micro-paths acts as a guide for cutting hypocotyls using a knife (Figure 1e, f). The detailed dimensions are shown in Figure S1. The lower stock part contains a seed pocket (width 900 μm), a root-growth path (length, 525 μm; width, 300 μm), and a root-growth space (length, 1.4 mm; height, 300 μm). The upper scion part contains a micro-path (length, 1.6 mm; width, 150 μm) with four pairs of pillars (diameter, 300 μm). The depth of these structures is approximately 450 μm, and the thickness of the scaffold is approximately 3 mm (Figure 1f). After sowing seeds onto the seed pockets, the micrografting chip is covered. The semi-solidified agar medium is retained in the region between the seed pocket and the root-growth space (Figure 1g).

**Figure 1.**
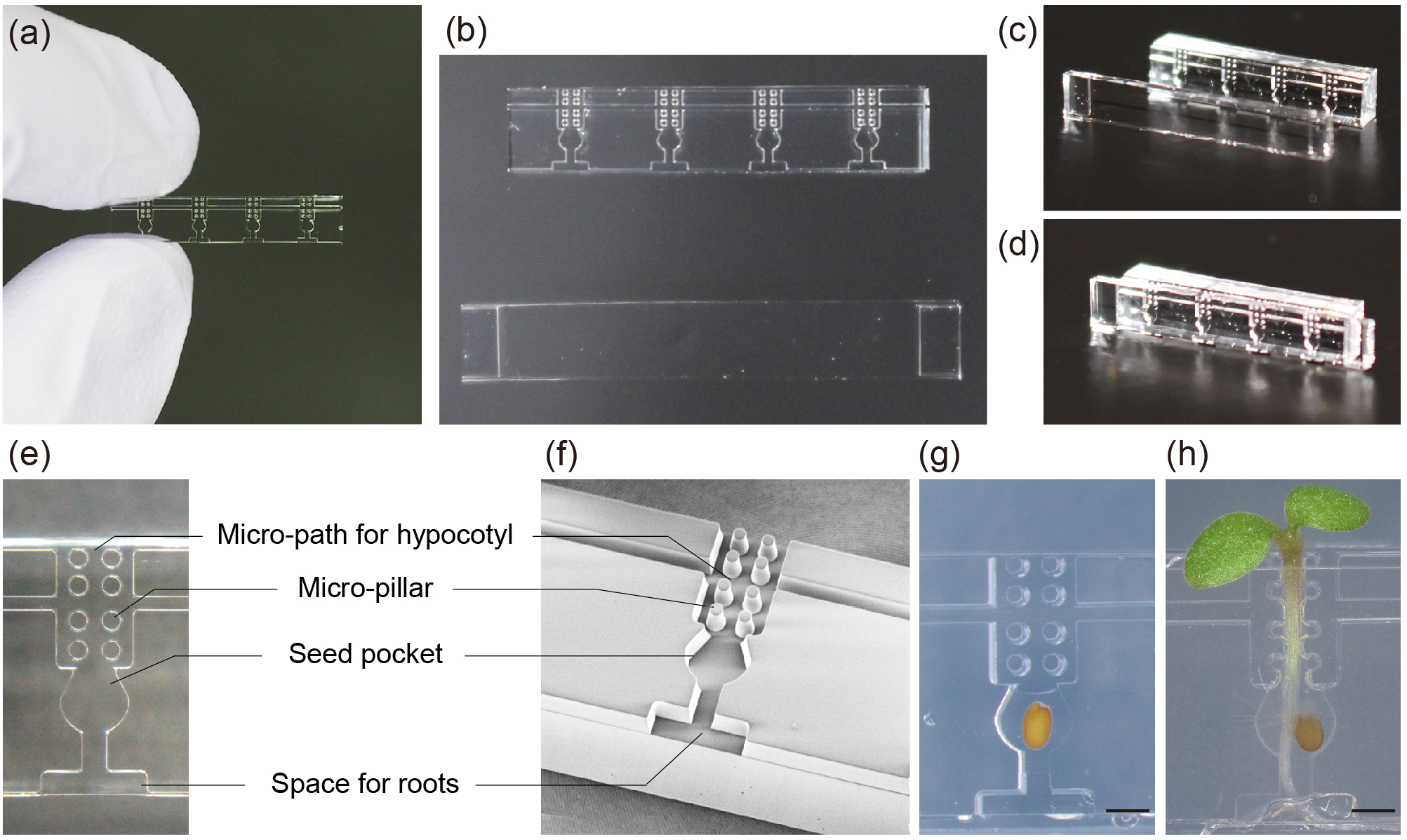
The structure of the micrografting chip. (a) The micrografting chip. (b) Image of a micrografting chip and its cover. (c, d) Images of a vertically oriented micrografting chip and its cover before assembly (c) and after assembly (d). (e) An enlarged image of a unit of a micrografting chip. (f) A scanning electron micrograph of a unit of a micrografting chip. (g) Image of a unit after sowing seeds. The seed pocket and space for roots are filled with seedling growth medium. (h) An image of a chip unit at 4 days after germination. Bars = 5 mm in (b) and 500 μm in (e), (f), (g), and (h).

Figure 1h shows a germinated seedling on the micrografting chip, the hypocotyl of which passed through the micro-path by pushing the pillars outward. Since the distance of the pillars, 150 μm, is narrower than the width of hypocotyls, mostly ranging from 200 to 250 μm, all seedlings encounter the pillars during their growth. The elastic PDMS pillars bend as the seedling becomes thicker, allowing passage of seedling hooks larger than the width of the micro-path. These pairs of pillars guarantee straight growth of the hypocotyl at the center of the micro-paths by pushing the hypocotyl inward. Because the pillars immobilize the seedling during grafting and the latter curing period, the accuracy of grafting manipulation is increased.

### Micrografting using the silicone chip

An outline of the micrografting using the chip is shown in Figure 2. The entire process takes 15 days from sowing the seeds on the chip to confirmation of successful micrografting. On day 1, pre-watered sterilized seeds were sown in the seed pockets of the micrografting chip in partly solidified 0.4% agar to retain water in the seed pocket spaces (Figure 1g). We often filled the root-growth space by semi-solidified agar as well, but filling only the seed pocket was sufficient for seed germination. Next, the chip was covered (Figure 3a) and placed vertically onto a membrane on half-strength Murashige– Skoog (MS) agar medium. By covering the chip, the growth of the seedlings is restricted to along the micro-paths. For the first 2 days, plants were grown in darkness (the plates were covered with aluminum foil) to make the seedlings etiolated. During this period, the hypocotyls elongated along the micro-paths without cotyledon opening, pushing the pillars outwards. On day 3, when the apical part of the hypocotyl–cotyledon hook reached the top of the micro-path, the aluminum foil cover was removed, and the plants were grown in light at 22°C for 2 days to promote greening of the shoots. Next, the cotyledons passed through the micro-path completely and fully expanded (Figure 3b). Finally, the plants were grown on the chip until evaluation of the grafting success (Figure 3c).

**Figure 2.**
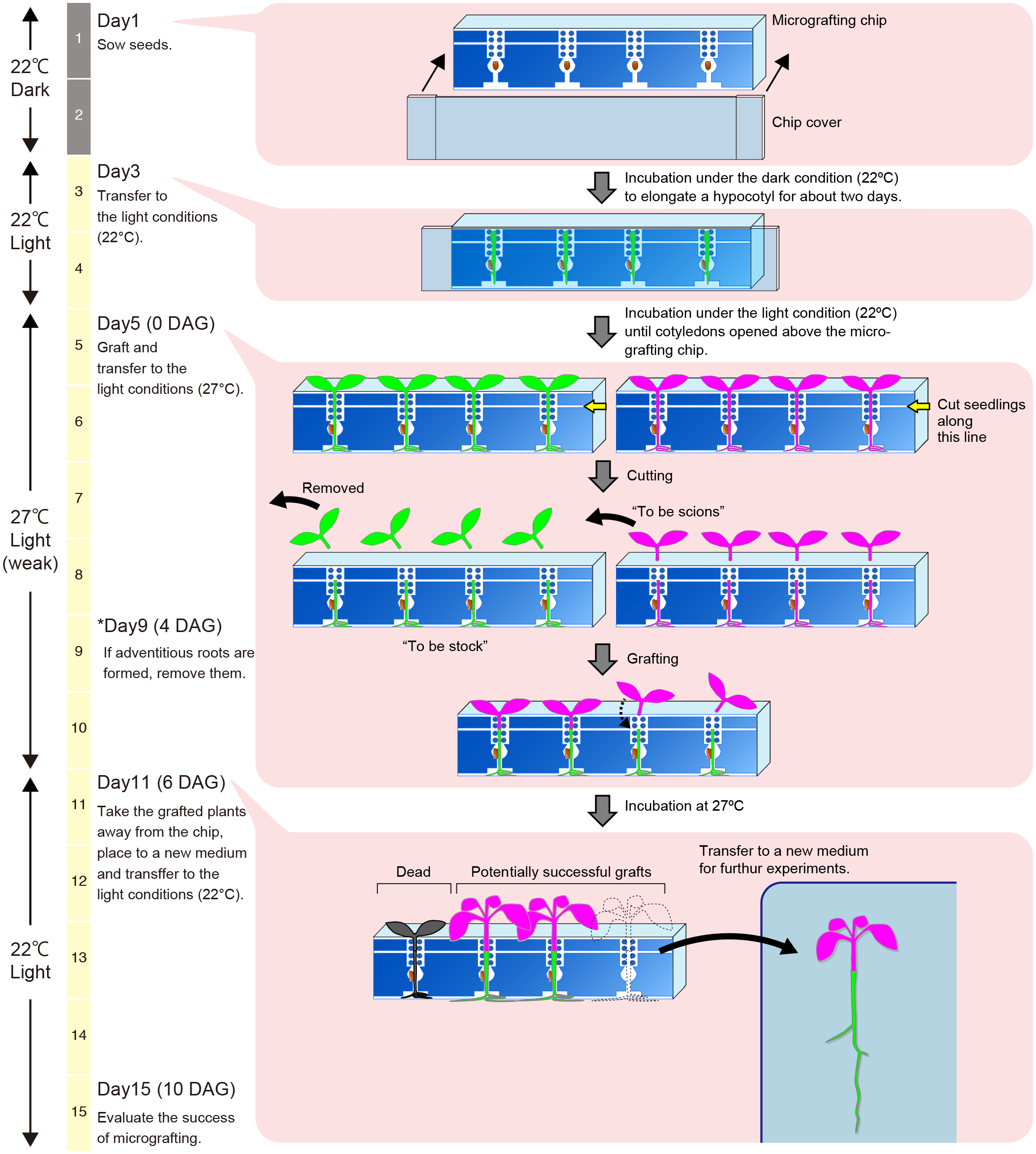
Micrografting using the micrografting chip. The micrografting procedure, with scion plants indicated in magenta and stock plants in green. The process takes 15 days from sowing seeds on the chip to confirmation of successful micrografting. On day 1, seeds were sown on the chip and placed in the dark. On day 3, plants were transferred to light conditions and grown for 2 days. On day 5 (0 days after grafting, 0 DAG), stock and scion plants were prepared and assembled on the chip to produce grafts. The plants were incubated at 27°C for 6 days. On day 11 (6 DAG), successful grafts were transferred to new medium. On day 15 (10 DAG), successful grafts had formed functional connections and thereafter were used for analysis.

**Figure 3.**
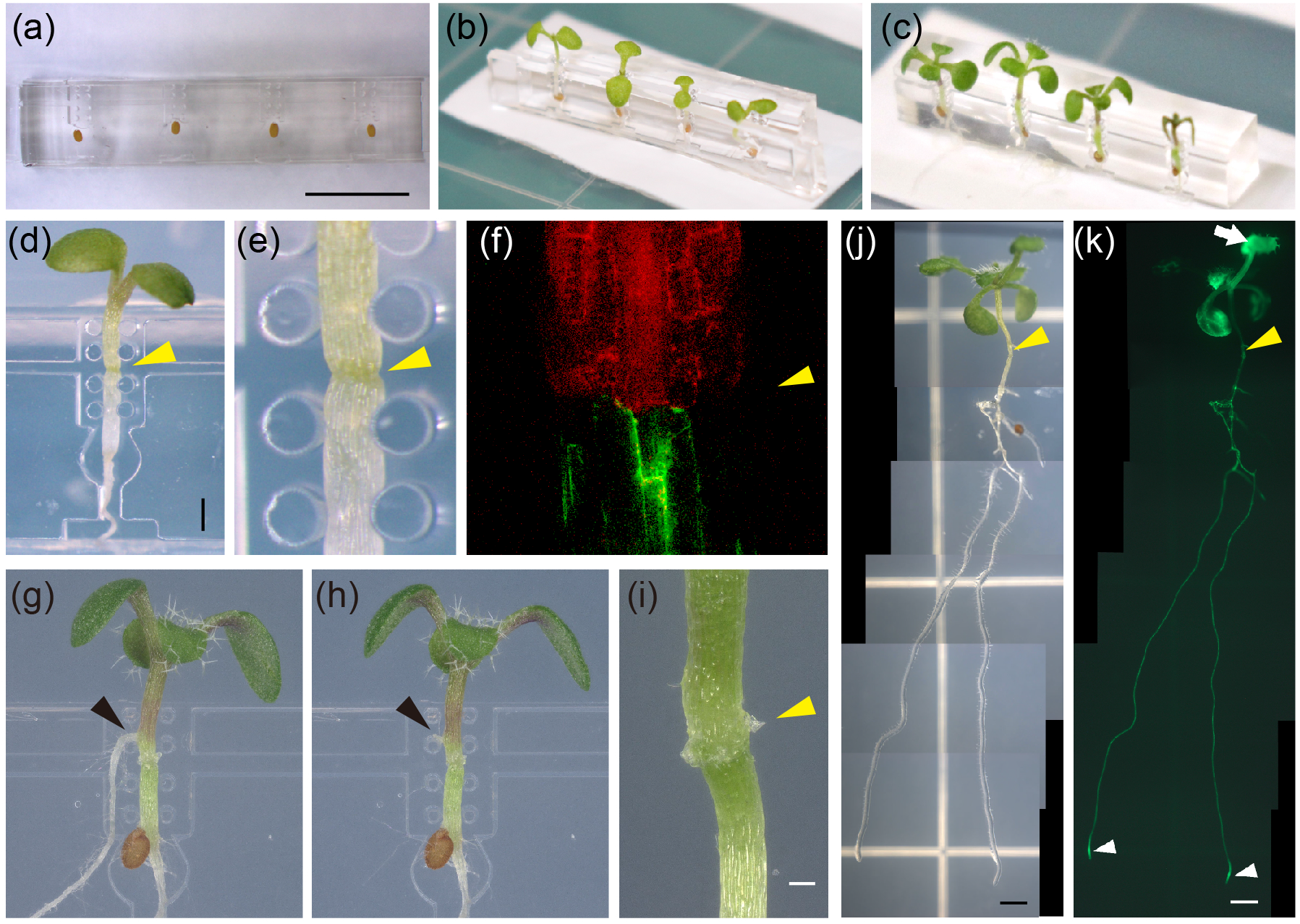
Grafted plants at the indicated time points during micrografting. (a) Image of a micrografting chip immediately after sowing seeds (day 1). (b) Image of a micrografting chip at 4 days after sowing (day 5). (c) Image of a micrografting chip at 6 days after grafting (day 11). Black arrowhead indicates a withered seedling. (d, e) A seedling immediately after micrografting (d) and an enlarged view of the grafting point (e). (f) A confocal image of a graft junction in the micrografting chip. The scion harbored *RPS5A::LTI:tdTomato* (red), and the stock harbored *35S::GFP* (green). (g) (h) A grafted seedling at 4 days after grafting (day 9). An adventitious root emerging from the scion hypocotyl (black arrowhead) was removed. (i) An image of a graft junction at 6 days after grafting. (j, k) Images of a grafted seedling at 10 days after grafting (day 15). CFDA was applied to a true leaf of the scion (white arrow), and CF fluorescence was imaged; bright field (j) and fluorescence (k) images. White arrowheads indicate CF fluorescence at the root tips of stock plants (k). Yellow arrowheads indicate graft junctions. Bars = 5 mm in (a), 500 μm in (d), (g), (h), and (i); 300 μm in (j); 100 μm in (e), (i), (j), and (k); and 50 μm in (f).

On day 5, micrografting was performed. Scion and stock plants were grown in separate micrografting chips to prevent mixing of materials. To perform micrografting, two chips (one each for the stock and scion plants) were transferred onto moist paper on a new plastic Petri dish. The chip covers were removed to expose the hypocotyls. We prepared and assembled one stock and one scion plant and performed grafting. For the stock plant, we cut the hypocotyl along the horizontal slot in the chip and removed the shoot. We next cut the hypocotyl of the scion plant in the same manner. The scion shoot part was transferred to the chip of the stock plant, and the scion was immediately assembled onto the stock plant using forceps. The locations of the scion and stock hypocotyls were adjusted using forceps such that the cut surfaces were in contact (Figure 3d and e). Grafting of two lines expressing green (GFP) or red fluorescent protein (RFP) showed the well-aligned scion–stock hypocotyls at the graft boundary (Figure 3f). After grafting, the chip was transferred onto 2% MS agar medium to reduce water evaporation, because excess moisture inhibits graft formation (Notaguchi *et al.*, 2009). The grafted plants were incubated in light at 27°C for 6 days. At 4 days after grafting, we checked the grafted plants; any adventitious roots on the scion hypocotyl were removed using fine scissors and forceps (Figure 3g and h). At 6 days after grafting (day 11), healing of the graft junction had progressed, and proliferated calli were observed at the graft junction (Figure 3i). At this time, successful grafts produced new true leaves, whereas unsuccessful grafts showed growth arrest (Figure 3c, asterisk). Successful grafts were transferred onto 1% MS agar and grown for 4 days for evaluation of grafting success on day 15.

Until 10 days after grafting (day 15), the grafted plants established vascular reconnections at the graft junction, and the growth of shoots and roots restarted. True leaves emerged on the scion, and primary root growth and lateral root formation were observed on the rootstock. We confirmed the establishment of phloem connections by visualizing transport of a symplasmic tracer dye, carboxyfluorescein (CF). Carboxyfluorescein diacetate (CFDA), the diacetate residue of which confers membrane permeability and which is digested by intracellular esterases to produce CF, was applied onto a scion leaf; CF fluorescence was detected in its rootstock, suggesting that CF was transported from the scion to the rootstock through the reconnected phloem tissues (Figure 3j, k). Also, the vigorous growth of grafted plants indicated successful graft formation. These results demonstrated that functional graft unions were established using the micrografting chip.

### Effects of environmental conditions on grafting

Although the presence of sugars in the medium increases the risk of fungal contamination, sugars also increase plant vigor and accelerate graft recovery (Marsch-Martínez *et al.*, 2013, Osugi *et al.*, 2017). To test the effect of sugar on the grafting success rate, we incubated grafted plants on MS agar containing 0% or 0.5% sucrose. We prepared a clean workplace using a clean air supply system (see Experimental Procedures) to minimize the risk of fungal contamination. The median success rate of grafting on non-sucrose medium was 35%, compared with 85% in medium containing 0.5% sucrose (Figure 4a). Therefore, a carbon source in the medium promoted graft reunion, and thus we used 0.5% sucrose in subsequent experiments.

**Figure 4.**
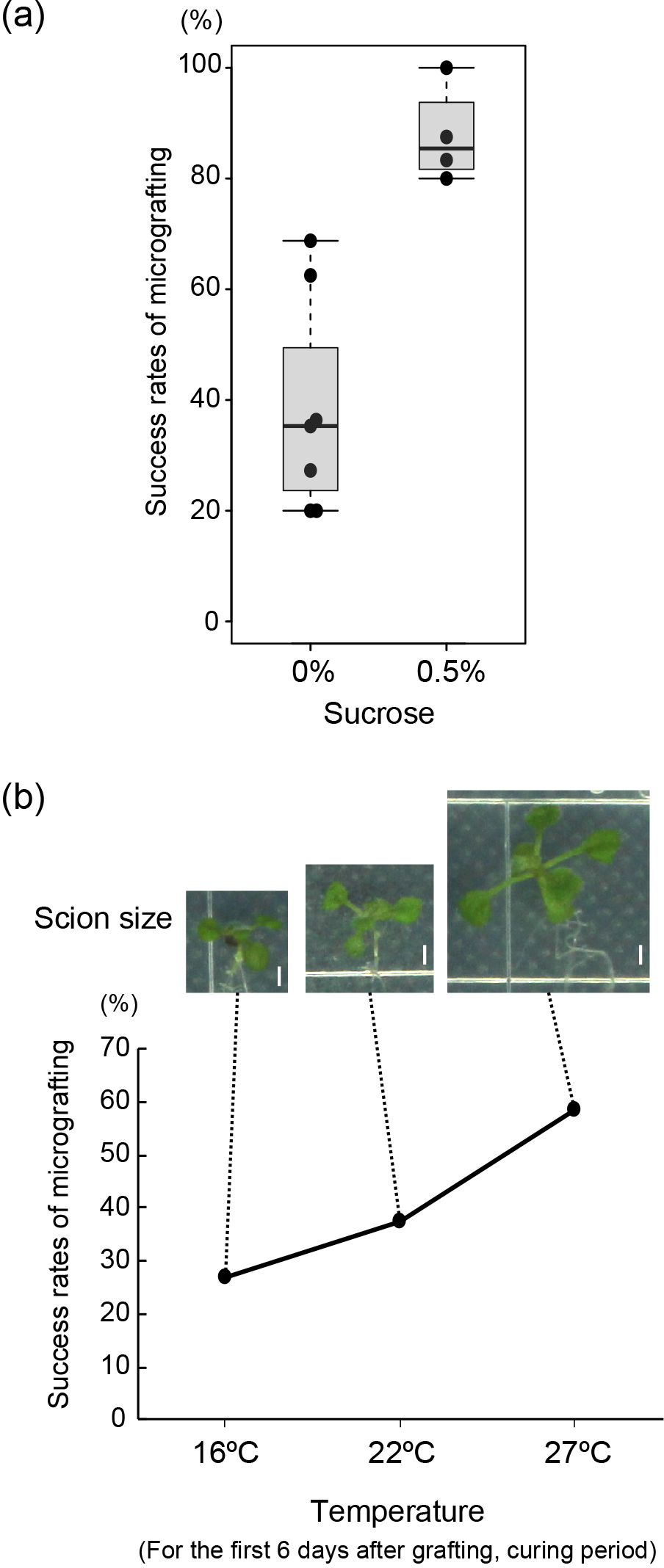
Effects of environmental factors on the success rate of micrografting. (a) Success rate of micrografting in the presence of 0% and 0.5% sucrose. (b) Success rate of micrografting according to temperature. Representative images of scions at 14 days after grafting are shown at top. Bars = 3 mm.

We next examined the effect of temperature during graft healing on the graft success rate by placing grafted plants at 16, 22, or 27°C for 6 days after grafting. Subsequently, the grafts were transferred to 22°C for 4 days, and grafting success was evaluated based on outgrowth. At 16, 22, and 27°C, the numbers of successful grafts were 12 of 45 (27%), 16 of 43 (37%), and 28 of 48 (58%), respectively (Figure 4b). In all grafts, phloem connection was confirmed by CF tracer experiments. The numbers of grafts in which CF fluorescence was detected in rootstock were 17 of 45 (38%), 25 of 43 (58%), and 36 of 48 (75%) at 16, 22, and 27°C, respectively. Thus, CF tracer transport was established. We excluded some grafts based on their appearance, such as those showing accumulation of anthocyanins or little regrowth. Note that the size of scion shoots was affected by temperature (Figure 4b). The scion shoots of grafts incubated at 27°C were larger than those incubated at lower temperatures, possibly reflecting the efficiency of tissue reunion at the graft junction. Thus, 27°C was the optimum temperature in terms of graft success rate and scion outgrowth, in agreement with previous reports (Turnbull *et al.*, 2002, Notaguchi *et al.*, 2009). We used 27°C and a 6-day incubation after grafting for the micrografting protocol. Under these optimum conditions, three operators achieved 48–88% grafting success rates using the chip (Table 1).

**Table 1.**
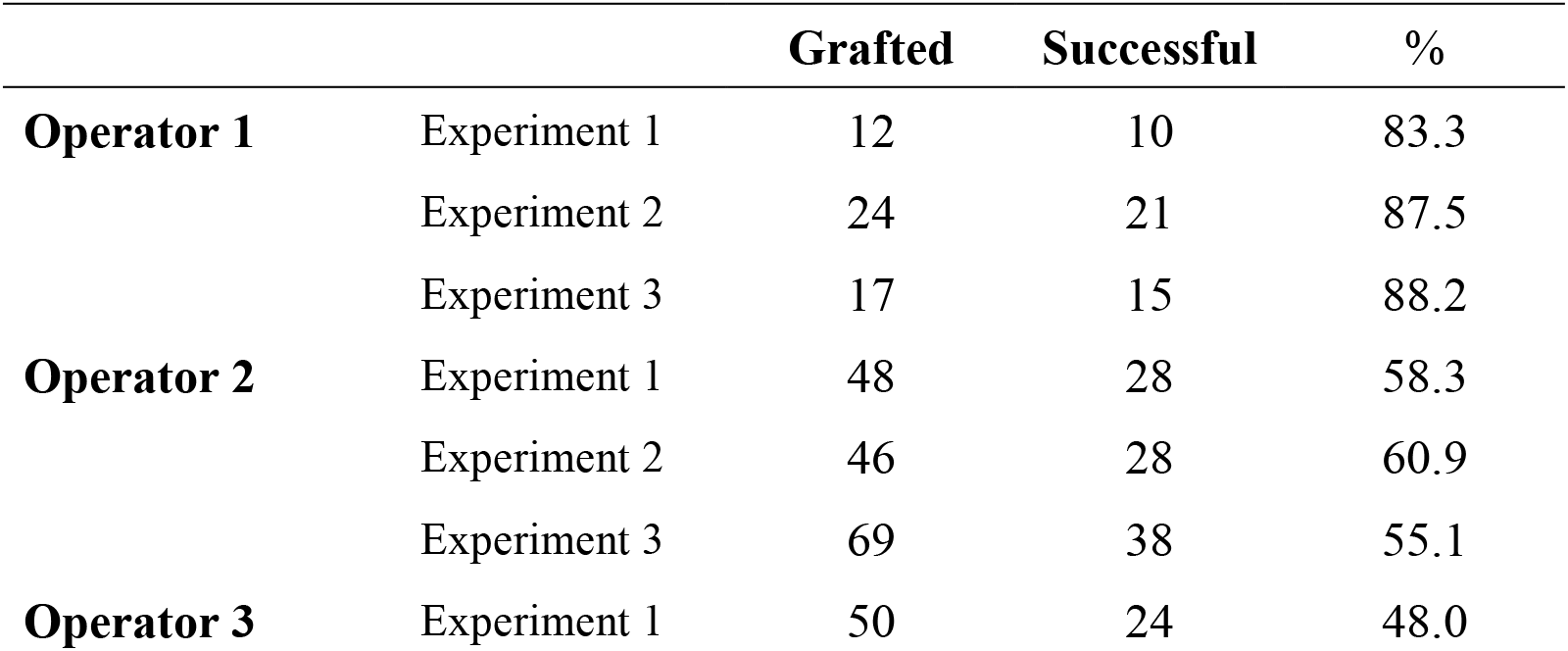
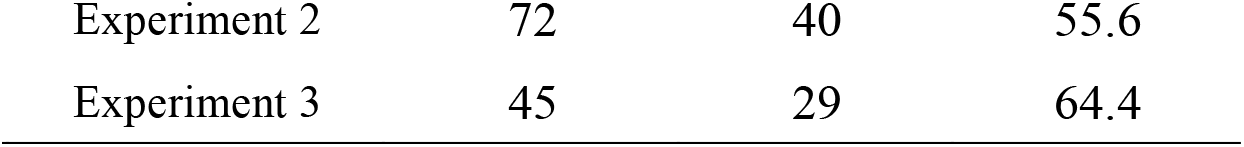
Success rate of grafting by three operators using the micrografting chip. Left to right: number of grafts, number of successful grafts 10 days after grafting, and grafting success rate (%).

|  |  | Grafted | Successful | % |
| --- | --- | --- | --- | --- |
| **Operator 1** | Experiment 1 | 12 | 10 | 83.3 |
|  | Experiment 2 | 24 | 21 | 87.5 |
|  | Experiment 3 | 17 | 15 | 88.2 |
| **Operator 2** | Experiment 1 | 48 | 28 | 58.3 |
|  | Experiment 2 | 46 | 28 | 60.9 |
|  | Experiment 3 | 69 | 38 | 55.1 |
| **Operator 3** | Experiment 1 | 50 | 24 | 48.0 |

|  |  |  |  |
| --- | --- | --- | --- |
| Experiment 2 | 72 | 40 | 55.6 |
| Experiment 3 | 45 | 29 | 64.4 |

### Evaluation of systemic control of iron translocation

We used the micrografting chip to characterize the role of NA in the systemic signaling regulating homeostasis of Fe, an essential micronutrient for plants and animals (Hänsch and Mendel, 2009). As a metal chelator, NA binds Fe and promotes its systemic transport (von Wiren *et al.*, 1999). However, the role of root- or shoot-derived NA in Fe ion transport was not examined independently. To investigate the contribution of Fe–NA transport in maintenance of Fe status at whole-plant level, we performed grafting using a quadruple mutant of the four Arabidopsis *NAS* genes, *nas4x-2*, which shows strong leaf chlorosis and a dwarf phenotype (Schuler *et al.*, 2012). Additionally, the *nas4x-2* mutant has defect in systemic signaling of Fe status resulting in the elevated expression of *iron-regulated transporter 1* (*IRT1*) in roots even under Fe-sufficient conditions (Klatte *et al.*, 2009). We conducted self and reciprocal grafting of WT and *nas4x-2* mutant plants using the micrografting chip. We first confirmed the chlorotic phenotype of the leaves of *nas4x-2* self-grafts, which was not observed in WT self-grafts. The shoots and roots of *nas4x-2* self-grafts were smaller than those of WT self-grafts. In *nas4x-2*/WT (scion/stock) grafts, in contrast, the chlorotic phenotype of the leaves of *nas4x-2* was rescued by grafting onto WT rootstock. Also, the *nas4x-2* scions were larger than those of *nas4x-2* self-grafts (Figure 5a). These observations indicate that, although it has been thought that NA is important for Fe transport in the phloem tissues in shoots (Jeong *et al.*, 2017), the lack of NA synthesis in the mutant shoot was complemented by the WT rootstock. The possible idea to explain this rescue is that NA itself is able to be transported from the WT rootstock to the mutant scion over graft junction. It is also possible that the root-to-shoot Fe–NA transport via xylem contributes in part of Fe shoot-ward translocation in Arabidopsis in addition to Fe–Ci conjugate, a major form of root-to-shoot Fe transport via xylem (Rellán-Álvarez *et al.*, 2010, Figure S2). The elevation of *IRT1* expression was not observed in the WT rootstock grafted with *nas4x-2* scion (Figure 5b), meaning that Fe status is correctly regulated in *nas4x-2*/WT grafted plants both in shoots and roots. Here, the restoration of Fe-deficiency response may be again reasoned by recruitment of NA from the WT rootstock to the *nas4x-2* mutant shoots, prior to Fe allocation from the mutant shoot to the entire body. In WT/*nas4x-2* grafts, both shoot and root grew normally (Figure 5a). This coincides with that Fe–Ci is a major form in Fe root-to-shoot transport and NA dependent transport of Fe is minor. The constitutive Fe-deficiency response in the roots of *nas4x-2* mutant was restored by grafting with WT scion shoots (Figure 5b). This result indicates that the regulation of Fe homeostasis is again regulated systemically and normal NA synthesis in the shoot is sufficient to control the systemic signaling for regulation of Fe absorption in roots. In WT/*nas4x-2* grafts, NA could redistribute from shoot to root in the form of Fe–NA conjugate and/or NA itself (Figure S2). Overall, it is indicated that not only shoot-derived NA but also root-derived NA contribute to Fe homeostasis in whole-plant level to achieve normal development of shoots and roots (Figure 5c).

**Figure 5.**
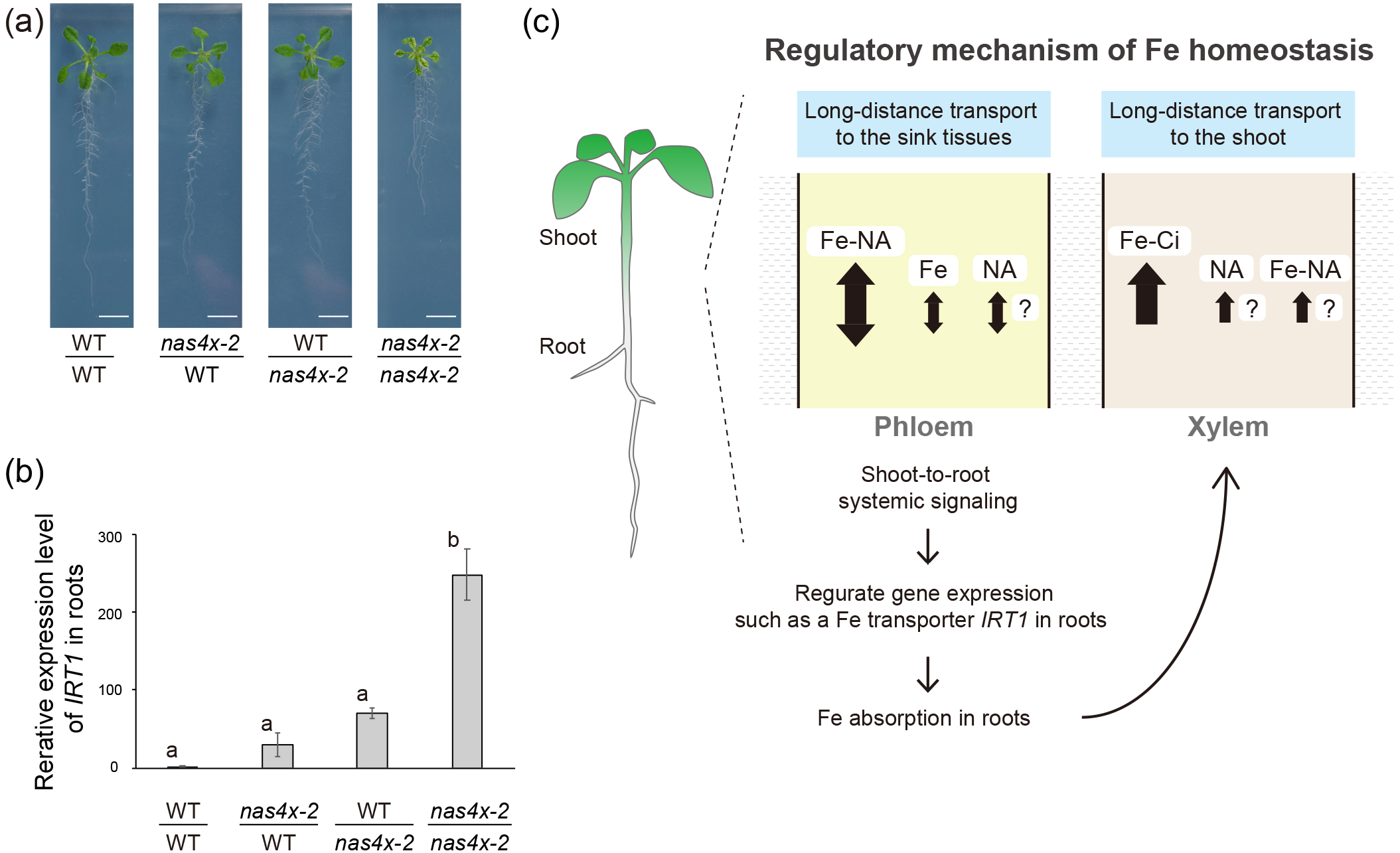
Micrografting of an NA synthase mutant of Arabidopsis using the micrografting chip. (a) Reciprocal micrografting of the WT and *nas4x-2* mutant. Images were obtained at 14 days after grafting. Bars = 1 cm. (b) qRT-PCR of *IRT1* expression in roots of grafted plants. Error bars indicate standard errors of the mean (WT/WT in quadruplicate and others in triplicate). Tukey’s test showed significant differences among the *nas4x-2*/*nas4x-2* grafts and the other graft combinations (*p*<0.01). (c) Model of systemic transport of Fe and its chelators, NA and Ci, between the shoot and root via the phloem and xylem. At the neutral pH in the phloem, the Fe^2+–^NA complex prevails. Na itself could be transported also. OPT3 transporter in shoots mediates the transport of Fe ion itself to the phloem. Thus, the transport of Fe ion in the other form(s) could happen in the phloem. In contrast, the Fe^3+–^Ci complex may be the major prevailing form of Fe in more acidic conditions in the xylem. This study indicates that NA itself and/or Fe–NA complex could ride on the xylem translocation stream. The shoot-to-root systemic signaling regulates Fe status by controlling the expression of Fe transporter gene, *IRT1*, in roots enhancing Fe absorption from the soil and Fe translocation to the shoot.

## Discussion

Recent studies on plant biology have focused on systemic physiology and gene function and have shown that signaling molecules are systemically mobile and coordinate plant growth and development (Liu and Chen, 2018, Ko and Helariutta, 2017, Tsutsui and Notaguchi, 2017, Okamoto *et al.*, 2016). The Arabidopsis micrografting method established by Turnbull *et al.* (2002) pioneered such research, and subsequent studies proposed modifications to their method (Notaguchi *et al.*, 2009, Yin *et al.*, 2012, Marsch-Martínez *et al.*, 2013). However, because of the small size of Arabidopsis seedlings, micrografting requires a high level of skill. To overcome this, we developed the micrografting chip based on MEMS technology (Figure 1 and Figure S1). This chip was used to prepare scion and stock plants and to perform all grafting procedures: seed germination, scion/stock assembly (grafting), and subsequent incubation until graft establishment (Figures 2 and 3). We optimized the procedure to obtain uniformly grafted plants (Figure 4). The success rate of micrografting using the chip was typically 48–88% (Table 1), similar to that for conventional micrografting in Arabidopsis (Turnbull *et al.*, 2002, Yin *et al.*, 2012, Marsch-Martínez *et al.*, 2013). Establishment of a functional vasculature was confirmed by observation of shoot growth (Figure 3c) and transport of CF (Figure 3k). Finally, we used the chip to investigate the function of *NAS* in systemic Fe transport using WT and mutant plants (Figure 5). Thus, our micrografting chip enabled investigation of systemically regulated plant biology.

The micrografting chip is made of PDMS, which has good biocompatibility, *i.e.*, low toxicity to plants. Also, the micro-path with elastic silicone pillars allows passage of cotyledons and holds the hypocotyl in place. Moreover, the micro-path with pillars was narrower than the width of hypocotyls, accepting a range of sizes of seedlings. Without elasticity of the micro-path, the shoot passing and the hypocotyl holding would not be accomplished concurrently. The flexibility of PDMS pillars was previously used to culture plant ovules, which also have size varieties and expand during their growth (Park *et al.*, 2014). Finally, the transparency of PDMS enabled observation of the grafting site (Figure 3f). The chip did not show autofluorescence at the graft junction (Figure 3f). Tissue reconstruction at the graft junction involves phytohormones, the wounding response, and vasculature development (Yin *et al.*, 2012, Matsuoka *et al.*, 2016, Melnyk *et al.*, 2018). Several fluorescent reporter lines for these phenomena are available, such as DR5 promoter-driven fluorescent proteins (Ottenschläger *et al.*, 2002) and the R2D2 reporter (Liao *et al.*, 2015) for the auxin response, *WIND1p::GFP* (Iwase *et al.*, 2011) for the wounding response, and *CALS7p::H2B:YFP* (Furuta *et al.*, 2014) and *SEOR1p::SEOR1:YFP* (Froelich *et al.*, 2011) for vasculature development. Our micrografting chip enables imaging of cell dynamics at the graft junction because the scion and stock plants remain attached during graft formation. The above features of the developed chip facilitate micrografting experiments.

We examined the effects of carbon source and incubation temperature on micrografting. Growth medium containing 0.5% sucrose increased the success rate of micrografting (Figure 4a). The cells surrounding the graft junction divide to fill the gap between the scion and the stock to complete grafting. Because glucose, a metabolite of sucrose, promotes cell proliferation (Wang and Ruan, 2013), it likely enhances graft establishment. Indeed, Marsch-Martínez *et al.* (2013) reported that sugar promoted graft recovery. An antibiotic, such as carbenicillin, can be added to the medium to reduce the risk of contamination. The optimum temperature for incubation after grafting was 27°C (Figure 4b), although that for Arabidopsis growth is 22–23°C (Rivero *et al.*, 2014). Gray *et al.* (1998) reported that high temperature increased the level of indole-3-acetic acid, the most common auxin, in Arabidopsis hypocotyls. Several auxin response genes, including *ALF4* and *AXR1*, are required for graft formation (Melnyk *et al.*, 2015). Therefore, incubation at a high temperature should promote graft formation via the auxin pathway. Adventitious root formation on the scion hypocotyl hampers micrografting. Before graft reunion, adventitious roots had formed on the scion hypocotyls because of the requirement for water (Figure 3g); we removed these roots (Figure 3h) to enhance the grafting success rate. The presence of adventitious roots suppresses grafting because scions can survive without graft union with rootstocks. Formation of adventitious roots is also triggered by auxin accumulation at the cut end of the scion hypocotyl (Yin *et al.*, 2012). Auxin is synthesized in leaves and translocated to roots, where it accumulates and induces cell division and sink establishment, leading to adventitious root formation (Druege *et al.*, 2016). However, auxin accumulation and subsequent induction of the expression of auxin-response genes are also required for graft formation (Melnyk *et al.*, 2015). To further increase the success rate of micrografting, the growth conditions and/or compounds that inhibit adventitious root formation without hampering graft formation need to be determined. Our micrografting chip will promote generation of uniform grafts and evaluation of the abovementioned conditions.

Using the micrografting chip, we conducted experiments to investigate systemic signaling for maintenance of Fe status. We performed reciprocal grafting of the WT and *nas4x-2* mutant, in which NA synthesis and Fe–NA transport were diminished, and showed that root- and shoot-derived NA contribute to Fe mobilization as well as systemic signaling to maintain Fe status in whole-plant level (Figure 5). The restoration of the *nas4x-2* root phenotype by WT shoot grafting emphasized the contribution of Fe–NA transport via shoot phloem tissues to systemic Fe distribution. Previous studies already showed that the Fe status is regulated both locally and systemically (Jeong *et al.*, 2017, Kumar *et al.*, 2017). With regard to local regulation, the amount of available Fe in the root apoplast affects the expression of *IRT1* and *FERRIC REDUCTION OXIGENASE2*, encoding a ferric chelate reductase (Vert *et al.*, 2003). Recently, Kumar *et al.* (2017) conducted micrografting experiments using a mutant of Fe–NA transporters called Yellow Stripe1-Like (YSL) family. A double-knockout mutant of two YSL members, *ysl1ysl3*, showed a weak Fe-deficiency response even under Fe-starvation conditions, the phenotype of which is opposite to the *opt3* and *nas4x-2* mutants. Kumar *et al.* (2017) indicated that the weak deficiency phenotype of *ysl1ysl3* mutant could be explained by the restriction of Fe distribution within leaf veins. The weak Fe-deficiency response of a *ysl1ysl3* mutant was rescued by grafting onto WT shoots but not WT roots, demonstrating that YSL1 and YSL3 in shoots, the expression of which is strongly detected in the xylem parenchyma and the phloem, respectively, are required for the Fe-deficiency response in roots. This suggested that Fe–NA transport possibly via xylem-to-phloem exchange is important to generate shoot-to-root systemic signals to maintain normal Fe status. In our grafting experiments using the *nas4x-2* mutant where the mutant was grafted as scion shoot or rootstock, the shoot-to-root systemic signal regulating Fe status was generated. Together with the previous research on YSL1 and YSL3, our data indicated the presence of Fe–NA conjugates in the shoot of both WT/*nas4x-2* and *nas4x-2*/WT grafts. If so, there are implications; (1) Fe is transported from the *nas4x-2* root or WT root to the WT shoot or *nas4x-2* shoot probably in a form of Fe–Ci conjugate or Fe itself and (2), in *nas4x-2*/WT grafts, the absence of NA synthesis partly in the shoot of *nas4x-2*/WT grafts was might restored by NA recruitment from the WT roots (Figure S2). Alternatively, there is other shoot-to-root systemic signaling pathway for Fe homeostasis independent of Fe– NA conjugate. Overall, the regulation of Fe uptake in roots was regulated by local and systemic Fe signaling in root-to-shoot-to-root recirculation of bioavailable Fe and NA.

We developed a micrografting chip for Arabidopsis and determined the incubation conditions that maximized the grafting success rate. Because eudicot seedlings have an identical body structure, the micrografting chip can be used for eudicot plants other than Arabidopsis, such as model plants and vegetables, by adjusting the size of the chip. The molds produced using an SU-8 photoresist and MEMS technology are < 500 μm in depth (Jin *et al.*, 2003); therefore, larger chips would require other mold-production techniques, such as three-dimensional printing or metalworking. Advances in analytical equipment (*e.g.*, high-throughput sequencers) and biotechnological tools (*e.g.*, genome-editing enzymes) have promoted plant biology research. Taken together with such progressing techniques, the micrografting chip will facilitate further studies on systemic signaling in Arabidopsis and possibly other plant species.

## Experimental Procedures

### Preparation of micrografting chips

The micrografting chips were designed using LayoutEditor software (Juspertor GmbH, Unterhaching, Germany). To prepare the mold, SU-8 100 (MicroChem, Westborough, USA) negative photoresist was stacked on a silicon wafer using a spin coater. The photoresist-stacked-wafer was covered with the photomask and exposed to ultraviolet light to cure the photoresist. Next, the wafer was washed with SU-8 developer (MicroChem) to remove uncured photoresist, and the mold was coated with trimethoxysilane (Sigma, St. Louis, USA). To produce the micrografting chip, a mixture of SILPOT184 PDMS prepolymer and its curing agent (Dow Corning Toray, Tokyo, Japan) (10:1 [w/w]) was poured onto the mold and baked for 90 min at 65°C. Next, the micrografting chip was peeled from the mold. Before use, the chip was immersed in 76.9– 81.4% (v/v) ethanol for 2 days and autoclaved. Scanning electron microscopy images were captured using a VHX-D510 microscope (Keyence, Osaka, Japan) following the manufacturer’s instructions.

### Plant materials

*Arabidopsis thaliana* (Col-0) plants were used in this study. The details of the fluorescent lines *RPS5A::LTI:tdtomato* and *35S::GFP* are provided in Mizuta *et al.* (2015) and Notaguchi *et al.* (2009), respectively.

### Micrografting using the micrografting chip

All procedures were performed in a clean work area generated using a KOACH filtered air supply system (Koken, Ltd., Tokyo, Japan). MS agar (half-strength MS salt, 0.05% 2-(N-morpholino)ethanesulfonic acid hydrate, 1% or 2% agar) containing 0.5% sucrose was prepared in square Petri dishes. An autoclaved nylon membrane, Hybond-N+ (GE Healthcare, Chicago, USA), was placed on the MS plate with 1% agar, and an autoclaved micrografting chip was placed atop the membrane. A sterilized seed in 0.4% agar was placed in a seed pocket of the micrografting chip, and the chip was covered, incubated vertically in the dark at 22°C for 48 h, and transferred to light at 22°C. After incubation for 2 days, the cover was removed, and a hypocotyl was cut into two parts along the slot in the chip. The scion and stock were assembled on the chip using micro-pillars, and the chip was placed on 2% MS agar and incubated in light at 27°C for 6 days. Four days after grafting, the adventitious roots were removed from the scion. Six days after grafting, the plant was transferred to fresh 1% MS agar and incubated in light at 22°C for 4 days. The fluorescence signal was detected with a laser scanning microscope LSM5 PASCAL (Zeiss, Oberkochen, Germany).

### CFDA staining

Ten days after grafting, a small cut was made on true leaves of the grafted plants using scissors, and 0.5 mg/mL 5(6)-Carboxyfluorescein diacetate (Sigma) was applied to the cut using a P2 pipette. After ~2 h, translocation of CFDA to the root was observed using Axio Zoom V16 (Zeiss) and Axiocam 506 color stereomicroscopes (Zeiss).

### Quantitative reverse-transcription polymerase chain reaction of *IRT1*

Total RNA extraction, cDNA preparation and quantitative reverse-transcription polymerase chain reaction (qRT-PCR) were conducted as described previously (Tabata *et al.*, 2014). Total RNA was isolated from the roots of grafted plants at 14 days after grafting and used to produce cDNA for qRT-PCR amplification. The sequences of the primers were as follows: EF1α-F, 3’-CTTGGTGTCAAGCAGATGATTT-5’; EF1α-R, 3’-CGTACCTAGCCTTGGAGTATTTG-5’; IRT1-F, 3’-GGGTCTTGGCGGTTGTATC-5’; and IRT1-R, 3’-ACGCCATAACAAATTTCTTCATATT-5’.

## Acknowledgements

We thank Y. Hakamada, M. Hattori, and R. Masuda for their technical assistance, M. Mori for providing grafting operation data and H. Shikata for the scientific advice. We are grateful to D. Kurihara for providing seeds of *RPS5A::LTI:tdtomato* and *35S::GFP* and to P. Bauer for seeds of the *nas4x-2* mutant. This work was supported by grants from the Japan Science and Technology Agency (ERATO JPMJER1004 to T.H. and START 15657559 and PRESTO 15665754 to M.N.), Grants-in-Aid for Scientific Research (18KT0040 and 18H04778), and a grant from the Cannon Foundation (R17-0070) to M.N.

## Author contributions

HT, NY, RT and MN conceived of the research and designed experiments. HT, NY, YK, SI, YS, RT, HA and MN performed experiments and analyzed data. HA, TH and MN supervised the experiments. HT, RT and MN wrote the paper.

## Supporting information

**Figure S1**. The dimensions of the micrografting chip and its cover. (a) Dimensions of the micrografting chip. (b) Dimensions of a chip cover. (c) Dimensions of a unit of the micrografting chip.

**Figure S2**. Models of iron status in the reciprocal grafts of WT and *nas4x-2* mutant plants. Self- or hetero-graft combinations of WT (green) and *nas4x-2* mutant (magenta) are illustrated. Hypothesized transport of nicotianamine (NA), Ci and Fe between the scion and stock plants via the phloem and xylem as well as the phenotype of the scion shoot and rootstock is shown for each graft combination.

