## Supplemental file for "Micrografting device for testing environmental conditions for grafting and systemic signaling in Arabidopsis"

(a)

(b)

(c)

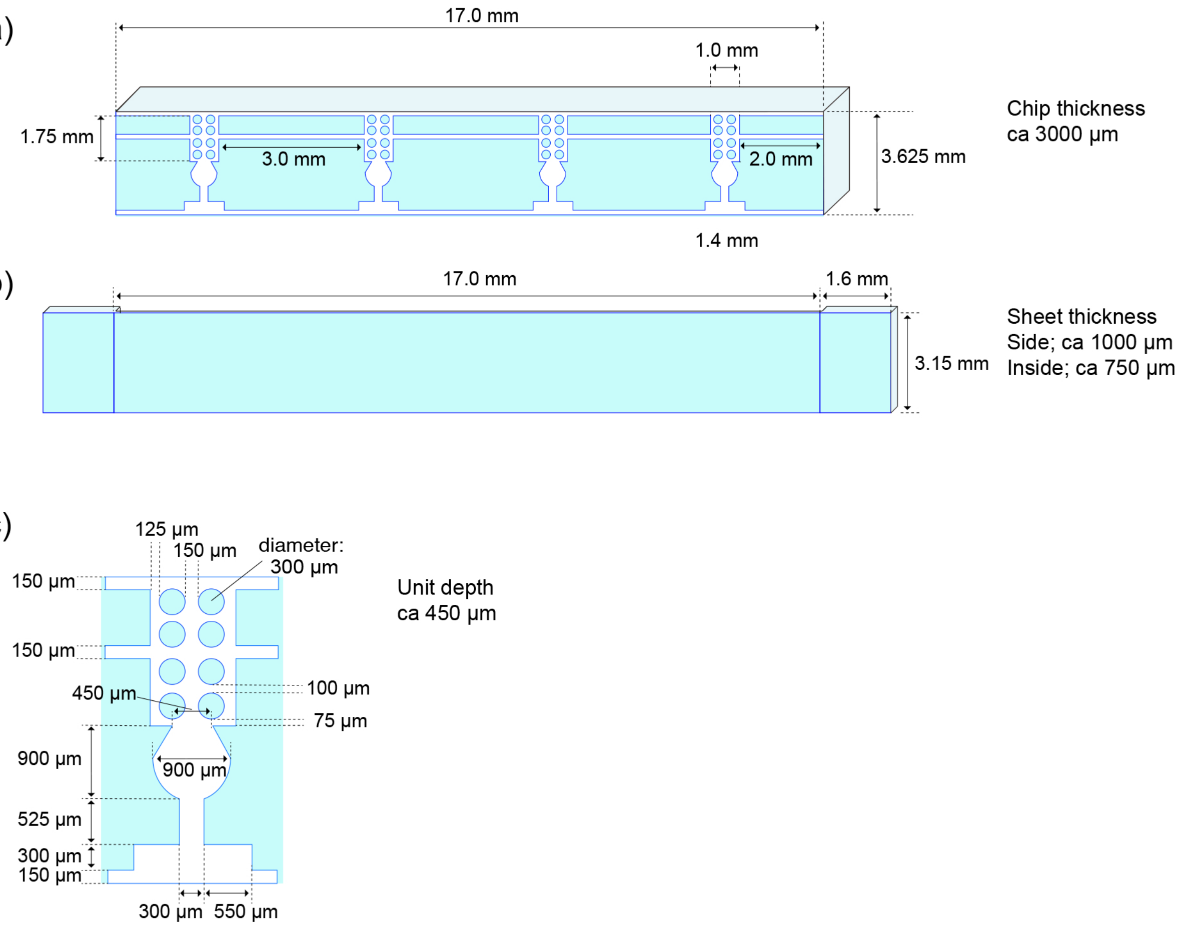


**Figure S1. The dimensions of the micrografting chip and its cover**
(a) Dimensions of the micrografting chip. (b) Dimensions of a chip cover. (c) Dimensions of a unit of the micrografting chip.

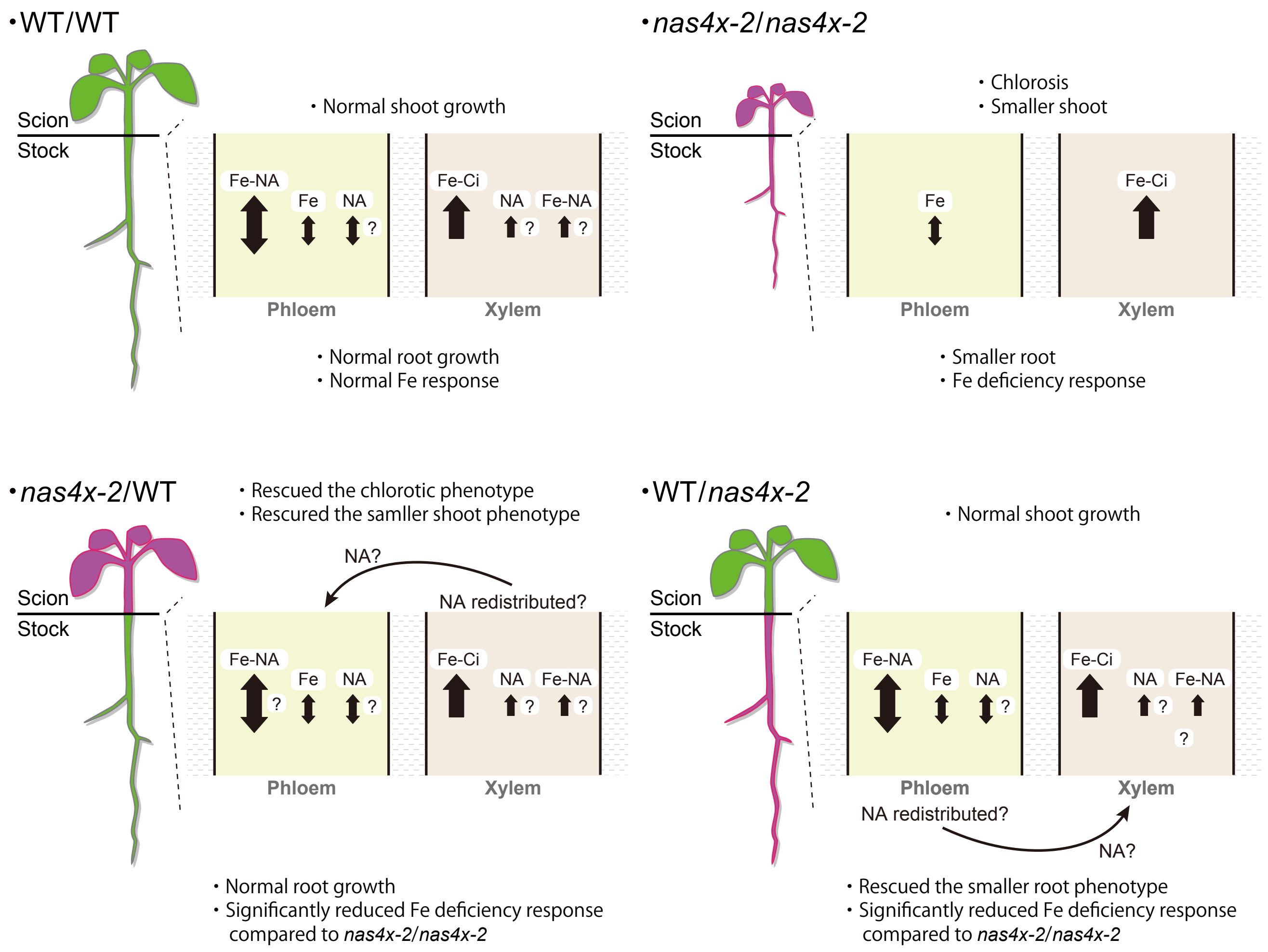


**Figure S2. Models of iron status in the reciprocal grafts of WT and nas4x-2 mutant plants**
Self- or hetero-graft combinations of WT (green) and nas4x-2 mutant (magenta) are illustrated. Hypothesized transport of nicotianamine (NA), Ci and Fe between the scion and stock plants via the phloem and xylem as well as the phenotype of the scion shoot and rootstock is shown for each graft combination.
